# Sucrose promotes etiolated stem branching through activation of cytokinin accumulation followed by vacuolar invertase activity

**DOI:** 10.1101/2020.01.08.897009

**Authors:** Bolaji Babajide Salam, Francois Barbier, Raz Danieli, Carmit Ziv, Lukáš Spíchal, Paula Teper-Bamnolker, Jiming Jiang, Naomi Ori, Christine Beveridge, Dani Eshel

## Abstract

The potato (*Solanum tuberosum* L.) tuber is a swollen stem. Sprouts growing from the tuber nodes represent dormancy release and loss of apical dominance. We recently identified sucrose as a key player in triggering potato stem branching. To decipher the mechanisms by which sucrose induces stem branching, we investigated the nature of the inducing molecule and the involvement of vacuolar invertase (VInv) and the plant hormone cytokinin (CK) in this process. Sucrose was more efficient at enhancing lateral bud burst and elongation than either of its hexose moieties (glucose and fructose), or a slowly metabolizable analog of sucrose (palatinose). Sucrose feeding induced expression of the sucrose transporter gene *SUT2*, followed by enhanced expression and activity of VInv in the lateral bud prior to its burst. We observed a reduction in the number of branches on stems of *VInv*-RNA interference lines during sucrose feeding, suggesting that sucrose breakdown is needed for lateral bud burst. Sucrose feeding led to increased CK content in the lateral bud base prior to bud burst. Inhibition of CK synthesis or perception inhibited the sucrose-induced bud burst, suggesting that sucrose induces stem branching through CK. Together, our results indicate that sucrose is transported to the bud, where it promotes bud burst by inducing CK accumulation and VInv activity.

## INTRODUCTION

In plants, the growing shoot apex inhibits the outgrowth of axillary buds further down the stem to control the number of branches. This phenomenon is referred to as apical dominance (Phillips, 1975; Ferguson and Beveridge, 2009; Wingler, 2017; Barbier et al., 2019). In response to decapitation, plants have evolved rapid long-distance signaling, involving sugars, to release axillary buds and replenish the plant with new growing shoot tips (Mason et al., 2014; Fichtner et al., 2017). Since the pattern of shoot branching may reflect the strength of the sugar sink, a clearer understanding of the regulatory mechanisms underlying shoot branching is expected to contribute to an increase in crop yields (Otori et al., 2017; Salam et al., 2017).

Shoot branching is controlled by complex interactions among hormones, nutrients, and environmental cues (Ongaro et al., 2008; Müller and Leyser, 2011; Leduc et al., 2014; Barbier et al., 2015b; Rameau et al., 2015; Roman et al., 2016; Fichtner et al., 2017; Le Moigne et al., 2018). Auxin, strigolactones and cytokinins (CKs) are the main plant hormones involved in the regulation of bud outgrowth, forming a systemic network that orchestrates this process (Ferguson & Beveridge, 2009). The classical view centers on the opinion that a bioactive form of the phytohormone auxin, which is produced in young leaves at the shoot apex (Cline, 1994; Ljung et al., 2001) and subsequently transported basipetally down the shoot in the polar auxin transport stream (Blakeslee et al., 2005), restricts the development of axillary buds (Sachs and Thimann, 1964; Cline, 1994; Bennett et al., 2016). Strigolactones inhibit shoot branching, as demonstrated by exogenous strigolactone application to the bud and by the strong branching phenotype displayed by strigolactone-synthesis and signaling mutants (Rameau et al., 2015). The fact that auxin upregulates strigolactone-biosynthesis genes in the stem suggests strigolactones’ involvement in mediating the branching inhibition by auxin (Saeed et al., 2017). Indeed, auxin requires strigolactones to inhibit bud outgrowth, since exogenous auxin is unable to fully repress decapitation-induced branching in strigolactone-deficient mutants (Arite et al., 2007; Beveridge et al., 2000). In contrast to auxin, a role for CKs in bud outgrowth emerged decades ago when direct CK applications onto dormant buds promoted bud outgrowth (Sachs and Thimann, 1967; Hartmann et al., 2011; Dun et al., 2012). Isopentenyltransferase enzymes control a rate-limiting step in CK biosynthesis, and transcript levels of genes encoding these enzymes are modified in response to auxin levels. Repression of CK-biosynthesis genes by auxin is well known (Miyawaki et al., 2004; Nordström et al., 2004; Tanaka et al., 2006).

Prior to the hormone and genetics era of plant biology research, the nutrient diversion theory of apical dominance was predominant (Wardlaw & Mortimer, 1970), involving the simple idea that bud outgrowth is inhibited by competition for resources (Kebrom, 2017; Barbier et al., 2019). The theory was narrowed down to sugar nutrients, proposing that apical dominance is maintained largely by sugar demand of the shoot tip, which limits the amount of sugar available to the axillary buds (Mason et al., 2014; Rameau et al., 2015). Sugars are a major source of carbon and energy produced by plants in an autotrophic fashion. In higher plants, three types of sugars accumulate to comparably high levels, namely the two monosaccharides glucose and fructose, and the disaccharide sucrose (Jung et al., 2015). From the site of their synthesis, sugars are partitioned to sink tissues in a controlled manner via the vascular system.

From a growth perspective, axillary buds are regarded as sink organs that are photosynthetically less active and need to import sugars to meet their metabolic demand and support their growth (Roitsch and Ehneß, 2000). A bud’s growth capacity is reflected in its sink strength, which represents its ability to acquire and use sugars. Therefore, to sustain its outgrowth, the bud has to compete for sugars, which constitute its main source of carbon and energy. Bud outgrowth occurs concomitantly with (i) starch reserve mobilization in stem tissues, mostly in perennial plants, (ii) high activity of sugar-metabolizing enzymes, and (iii) increased sugar absorption in the bud (reviewed by Rameau et al. 2015). The role of sugar as an early signal triggering bud activity has been recently suggested. Mason et al. (2014) showed that sugar initiates rapid outgrowth of the basal bud in pea after shoot decapitation. A strong correlation between sugar availability and branching has also been observed in studies involving defoliation (Alam et al., 2014; Kebrom and Mullet, 2015), enhanced CO_2_supply (Burnett et al., 2016; Otori et al., 2017), and inhibition of sucrose degradation (Salam et al., 2017). Fichtner et al. (2017) demonstrated that changes in the level of bud trehalose 6-phosphate—a signal of sucrose availability in plants—are correlated with initiation of bud outgrowth following decapitation, suggesting that trehalose 6-phosphate is involved in the release of bud dormancy by sucrose. In addition, the onset of bud outgrowth in various species is tightly correlated with the expression of genes involved in sugar transport, metabolism and signaling (Chao et al., 2016; Girault et al., 2010; Rabot et al., 2012). These findings support the theory that the growing shoot tip inhibits bud outgrowth by being a strong sink for sugars, thereby depriving the axillary buds (reviewed in [Barbier et al., 2015b]).

A more direct and genetic underpinning of the sucrose connection to bud outgrowth has been achieved through studies of vacuolar invertase (VInv). Salam et al. (2017) showed that silencing *VInv* in potatoes results in excess sucrose availability and a branching phenotype for the potato tuber. Meristem-specific overexpression of cell-wall or cytosolic invertase in *Arabidopsis* changes the shoot branching pattern in a complex manner, differentially affecting the formation of axillary inflorescences, branching of the main inflorescence, and branching of side inflorescences (Heyer et al., 2004; Wingler, 2017). This suggests that the sucrose-to-hexose ratio affects stem branching pattern and might differentially interact with hormones associated with bud growth.

Sugar and hormone networks interact to regulate different developmental processes (Ljung et al., 2015). However, the mechanism underlying these interactions is not fully understood. Sucrose has been shown to strongly induce CK synthesis in *in-vitro* grown single nodes in rose, suggesting that this hormone might mediate the sucrose effect (F. Barbier et al., 2015). However, replacing sucrose with CK in the growth medium was not enough to trigger bud outgrowth from the rose nodes, suggesting that sucrose also triggers a pathway independent of CKs, or that a minimal amount of sucrose is required for CKs to promote bud outgrowth, or both (F. Barbier et al., 2015). In rose, light controls the sugar supply to the axillary buds (Girault et al., 2010). However, in contrast to CKs, sugar supply is unable to restore the decreased branching phenotype triggered by darkness or low light intensity (Roman et al., 2016; Rabot et al., 2012).

Since potato sprouts can grow in the dark, the potato tuber and sprouts serves as an ideal model to study shoot branching under conditions in which most of the sugars are fed exogenously without the intervention of photosynthetic products. Our previous study showed that an exogenous supply of sucrose, glucose, or fructose solution to detached etiolated sprouts induces their branching in a dose-responsive manner (Salam et al., 2017). Although an increase in sucrose level was observed in tuber parenchyma upon branching induction, sugar analysis of grafted stems showed no distinct differences in sugar levels between branching and non-branching scions. Furthermore, silencing of the VInv-encoding gene led to increased sucrose levels and branching of the tuber (Salam et al., 2017). The objective of the present study was to decipher the mechanism by which sucrose modulates bud burst and elongation in the etiolated sprout. We found that sucrose is more efficient than glucose and fructose together, or the slowly metabolizable sucrose analog palatinose, at enhancing lateral bud burst and elongation in etiolated sprouts. Sucrose feeding to the isolated stem-bud system led to increased CK content and VInv activity in the lateral bud base in association with bud burst and elongation. Furthermore, inhibition of CK synthesis or perception inhibited the sucrose-induced bud burst, suggesting that sucrose induces stem branching largely through CK.

## MATERIALS AND METHODS

### Plant material and storage conditions

Freshly harvested tubers of potato (*Solanum tuberosum* ‘Desiree’) were obtained from a potato-growing field in the northern Negev, Israel, stored at 14°C for 3 weeks for curing, and transferred to 4°C until use. Wild-type (WT) ‘Russet Burbank’ (RBK) potato tubers and *VInv*-silenced lines RBK1, RBK22, RBK27 and RBK46 (Zhu et al., 2014) were grown in a greenhouse with controlled atmosphere (10 h day length at a temperature of 18–22°C; extra light was supplied by lamps with intensity ranging from 600 to 1000 μmol m^−2^ s^−1^). Water and mineral nutrients were provided by subirrigation for 5 min day^−1^. For sprouting induction, tubers were transferred from 4°C to 14°C, both under dark conditions. In all experiments, sprouts with three nodes were selected unless otherwise stated. Tubers and sprouts in all treatments were maintained at 95% relative humidity.

### Exogenous application of sugars, CK and CK inhibitors

Sprouts were detached manually from tubers stored at 14°C and surface cleaned by washing with sterile water for 5 min. Sprouts were dried for 3 min on a filter paper, and placed in sterile Eppendorf rack containing 300 mM sucrose, sorbitol, palatinose or a mix of glucose and fructose (300 mM each). They were then incubated at 14°C in the dark for up to 16 days, unless otherwise stated.

To evaluate the effects of CK on bud outgrowth, the synthetic CK 6-benzylaminopurine (BAP; Duchefa, Netherlands), as well as the CK-synthesis inhibitor lovastatin (Sigma, Israel) and CK-perception inhibitors LGR-991 and PI-55 (Spíchal et al., 2009; Nisler et al., 2010) were exogenously supplied to sprouts at 200 μM, unless otherwise stated. Bud length was measured with a millimeter-scale held perpendicular to the stem. Branches were defined as lateral buds longer than 0.2 cm.

### RNA extraction and cDNA synthesis

The lateral bud located at the third node from the apical bud was sampled from five sprouts per replicate and immediately frozen at −80°C. The buds were ground and RNA was extracted according to Chen et al. (2015) with slight modifications. The powdered tissue was added to 800 μl pre-warmed (65°C) extraction buffer (100 mM Tris–HCl, pH 8.0, 25.0 mM EDTA, 2.0 M NaCl, 3% w/v cetyl-trimethylammonium bromide, 4% w/v polyvinylpyrrolidone 40, 3% w/v β-mercaptoethanol) and incubated for 45 min at 65°C. Chloroform:isoamylalcohol (24:1, v/v) was added when the mixture had cooled to room temperature. The mixture, in centrifuge tubes, was allowed to stand for 10 min and then centrifuged at 12,400*g* for 20 min at 4°C. The above steps were repeated. RNA was precipitated by the addition of 2 ml LiCl at a final concentration of 3.0 M and incubation for 2 h at −20°C. Following another centrifugation at 12,400*g*, 4°C for 20 min, the pellet was washed twice with a volume 2 ml of 70% ethanol, centrifuged for 10 min, and air-dried at room temperature. Finally, the pellet was suspended in 1% DEPC-treated H_2_O. The quality and quantity of the extracted RNA were respectively assessed by spectrometer (Thermo NanoDrop 2000, USA). DNA was removed by incubating the RNA with DNase (Invitrogen, USA) for 10 min at 37°C (1 μl DNase for 10 μg RNA). The reaction was stopped by adding DNase-deactivation buffer (Invitrogen) and incubating for 5 min at 70°C. cDNA was obtained by reverse transcription performed on 400 ng of RNA using reverse transcriptase (PCR Biosystems, USA).

### Gene-expression analyses

Quantitative real-time PCR (qRT-PCR) was performed with SYBR Green mix (Thermo Fisher Scientific, USA) using cDNA as a template, with the following program: 2 min at 50°C, 10 min at 95°C, then 40 cycles of 15 s at 95°C and 60 s at 60°C. The primers used for the qRT-PCR are given in Supplementary Table S1A. Specific sets of primers were selected according to their melting curves. Fluorescence detection was performed using a Step One Plus Real-Time PCR system (Applied Biosytems, USA). Quantification of relative gene expression was normalized using *Ef1α* expression as an internal control (Nicot et al., 2005).

### Enzyme extraction and activity

VInv activity was measured as described previously (Miron & Schaffer, 1991), with minor modifications. Nodal stems carrying buds (250 mg fresh weight [FW]) were ground in liquid nitrogen and subsequently dissolved in 1 ml extraction buffer containing 25 mM HEPES– NaOH, 7 mM MgCl_2_, 0.5 mM EDTA, 3 mM dithiothreitol, and 2 mM diethyldithiocarbamic acid, pH 7.5. After centrifugation at 18,000*g* for 30 min, the supernatant was dialyzed overnight against 25 mM HEPES–NaOH and 0.25 mM EDTA, pH 7.5, and used as a crude extract. VInv activity was measured by incubating 0.3 ml of 0.1 M citrate/phosphate buffer (pH 5.0), 0.1 ml crude extract and 0.1 ml of 0.1 M sucrose. After 30 min incubation at 37°C, glucose liberated from the hydrolysis of sucrose was quantified by adding 500 μl Sumner’s reagent (3,5-dinitrosalicylic acid) and immediately transferring the sample to heating at 100°C for 10 min to terminate the reaction, then chilling at 4°C (Sumner and Graham, 1921). The reduction of dinitrosalicylic acid to 3-amino-5-nitrosalicylic acid by glucose was measured by absorbance at 550 nm in a spectrophotometer (Amersham Biosciences, UK). Quantitation of glucose in each sample was based on glucose standards. VInv activity was expressed as nanomoles glucose formed per gram FW per minute.

### Translocation and accumulation of labeled sugars

To determine sugar translocation and accumulation, sprouts were detached and cut at the base to expose the vascular tissues, and then incubated in 1 μCi of [U-^14^C]sucrose or a mixture of [U-^14^C]glucose + [U-^14^C]fructose, to a depth of 1 cm, supplemented with a mixture of 100 mM glucose and fructose, or sucrose. Sprouts were fed for 2 or 4 h. A total of 100 mg tissue was subsequently collected from the node, five buds per replicate. Radioactive counts of sucrose and the glucose–fructose mixture were determined by liquid-scintillation counting after crushing the tissue and diluting in Ultima Gold liquid scintillation cocktail (PerkinElmer, Israel) using a Packard Tri-Carb 2100TR counter analyzer (Packard BioScience, USA).

### Analysis of CK content

For each sample, 200 mg of freeze-dried powder of tissue was extracted with 1 ml of isopropanol:methanol:glacial acetic acid (79:20:1, v/v), and two stably labeled isotopes were used as internal standards and added as follows: 1 ng of [^15^N]*trans*-zeatin, 1 ng of [^2^H_5_]*trans*-zeatin riboside. The extract was vigorously shaken for 60 min at 4°C in a Thermomixer (Eppendorf), and then centrifuged (14000 *g*, 4°C, 15 min). The supernatants were collected, and the pellets were re-extracted twice with 0.5 ml of the same extraction solution, then vigorously shaken (1 min). After centrifugation, the three supernatants were pooled and dried (final volume 1.5 ml). Each dry extract was dissolved in 2000 μl of methanol:water (50:50, v/v), filtered, and analyzed by UPLC-Triple Quadrupole-MS (Waters Xevo TQ MS, USA). Separation was performed in a Waters Acquity UPLC BEH C18 1.7 μm 2.1 x 100 mm column with a VanGuard precolumn (BEH C18 1.7 μm 2.1 x 5 mm). Chromatographic and MS parameters for the CK analysis were as follows: the mobile phase consisted of water (phase A) and acetonitrile (phase B), both containing 0.1% formic acid in gradient-elution mode. The solvent gradient was applied as follows [*t* (min), % A]: (0.5, 95%), (14, 50%), (15, 5%), (18, 5%), (19, 95%), (22, 95%); [*t* (min), % B]: (0.5, 95%), (14, 50%), (15, 5%), (18, 5%), (19, 95%), (22, 95%); flow rate was 0.3 ml min^−1^, and column temperature was kept at 35°C. CK analyses were performed using the ESI source in positive ion mode with the following settings: capillary voltage 3.1 kV, cone voltage 30 V, desolvation temperature 400°C, desolvation gas flow 565 l h^−1^, source temperature 140°C. The parameters used for multiple reaction monitoring (MRM) quantification of the different hormones are shown in Table S1B.

### Data analysis

Data were analyzed using Microsoft Excel 2010. ANOVA and Tukey–Kramer test were performed using JMP software (version 3 for windows; SAS Institute).

## RESULTS

### Sucrose induces lateral bud elongation better than hexoses

We recently showed that sucrose and its hydrolytic products induce stem branching in a dose-responsive manner under etiolated conditions (Salam et al., 2017). To distinguish between the effects of sugars on bud burst vs. bud elongation, we conducted a detailed time course of the differential effect of sucrose or a mix of glucose and fructose (hexoses) on the number of branches and lateral bud elongation. Tubers were incubated at 14°C until sprouting, and sprouts with three nodes were then detached manually, placed in 300 mM sucrose, hexoses (glucose + fructose, 300 mM each), sorbitol (an osmotic control) or water, and incubated at 14°C for 9 days. The effect of sucrose was also compared to palatinose, a sucrose analog (glucose‐1,6‐fructose) which is not imported into the cell and is only slowly metabolized by vacuolar invertase and sucrose synthase (Loreti et al., 2000; Sinha et al., 2002;Wu & Birch, 2010). Sucrose and hexoses induced branching and lateral bud elongation (Fig. 1). Water and the other control and sugar-related treatments had little or no growth effect. The sugar alcohol and osmotic agent sorbitol, which can be imported but is not generally (or only slowly) metabolized by plant cells (see Klepek et al., 2005), and the sucrose analog palatinose, were unable to induce branching and elongation (Fig. 1). Sucrose and hexoses yielded similar branching, but lateral bud elongation was significantly higher during the 9 days of sucrose vs. hexose feeding. These results suggest that sucrose and its hydrolytic products enhance both stem branching and elongation under etiolated conditions, with a significantly higher effect of sucrose on bud elongation. Thus sucrose, as a whole molecule, may have an advantage over hexoses or its analogs.

**Fig. 1.**
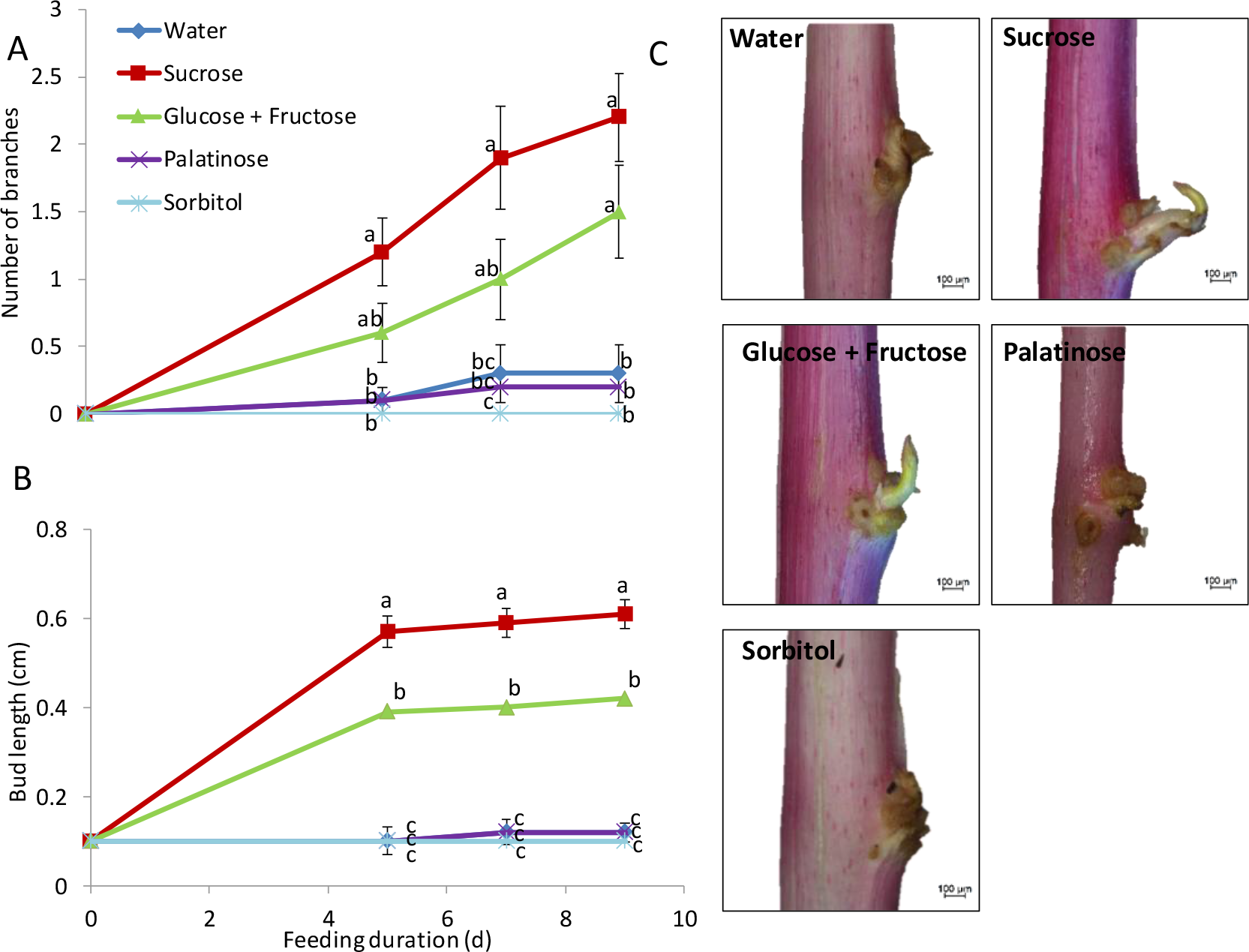
Exogenous sucrose or hexoses induce lateral bud burst and elongation in etiolated stems. Sprouts were detached from the tubers and supplemented with sugars (sucrose, glucose + fructose, palatinose, sorbitol, each at 300 mM) or water for 9 days at 14°C, 95% relative humidity, in the dark. **A**, Number of branches and **B**, lateral bud length, **C**, Images showing the lateral node after 7 days of treatment. Bars = 100 μm. Data represent averages of three experiments, each performed with 10 replicates per treatment. Error bars represent SE. Different letters represent significant differences between treatments at each time point (*P* < 0.05).

### Sucrose and hexoses translocate to the stem and penetrate the lateral bud

We previously reported that labeled sugars can be transported to the apical bud and lateral node of the etiolated stem following exogenous feeding of tuber parenchyma (Salam et al., 2017). To test whether sucrose and hexoses are translocated into the lateral bud itself or only to its base (node), we fed labeled sugars ([U-^14^C]sucrose, or [U-^14^C]glucose and [U-^14^C]fructose) to the base of detached stems (using the same system as above). After 2 h incubation with either sucrose or hexoses, we detected radioactivity at the node and inside the lateral bud (Fig. 2). Levels of radiolabel were unchanged in the node and lateral bud between 2 and 4 h of incubation (Fig. 2). While these results indicated translocation and entry into the lateral bud, it was not possible to distinguish whether the radioactivity measured in the buds was due to the movement of glucose and fructose, or to their reconversion to sucrose.

**Fig. 2.**
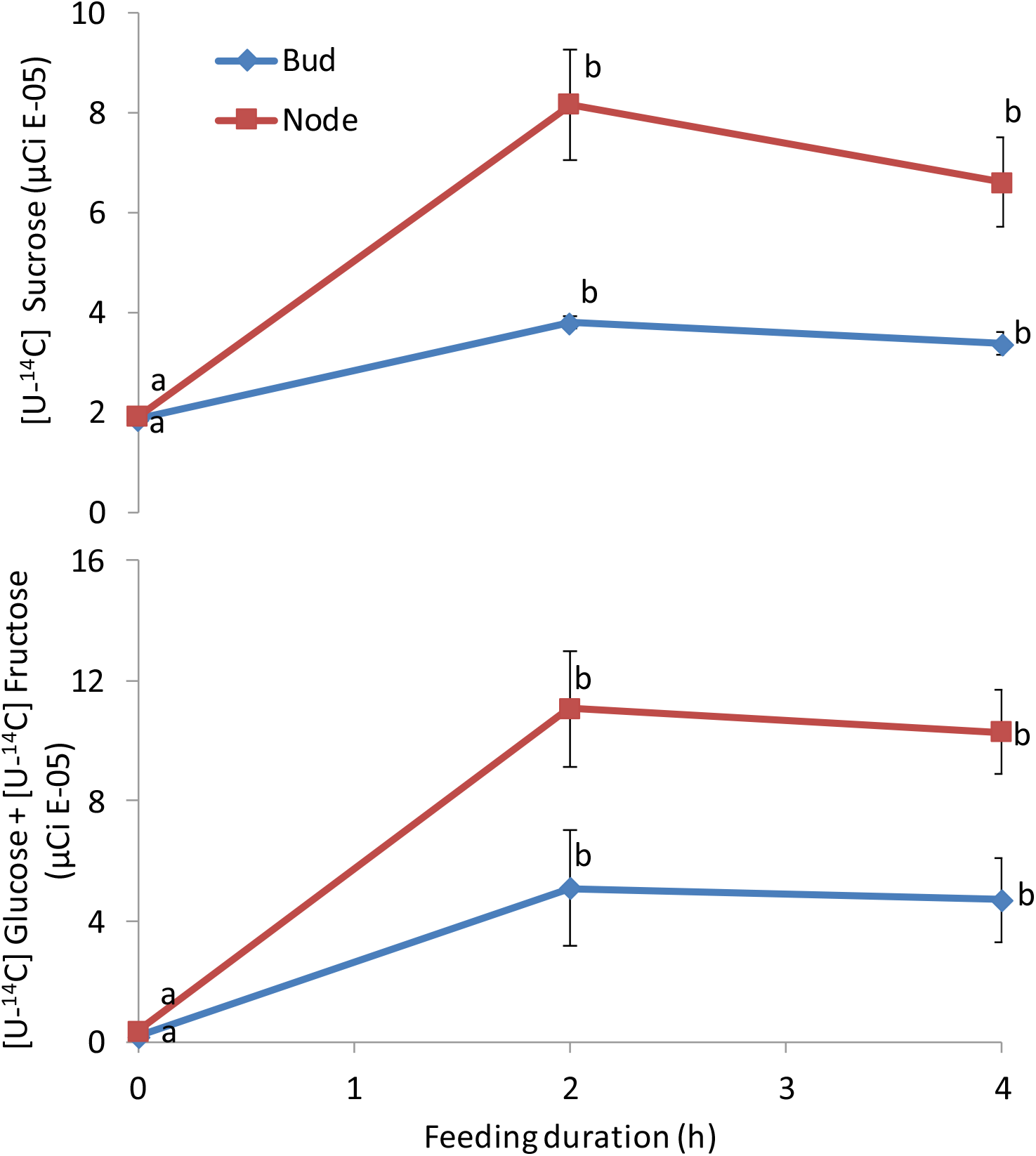
Sucrose and hexoses translocate from the stem into the lateral bud. Detached etiolated stems were fed with water solution containing **A**, 1 μCi [U-^14^C]sucrose or **B**, 1 μCi [U-^14^C]glucose + [U-^14^C]fructose for 0, 2 and 4 h in the dark. Each value is the mean of five independent measurements ± SE. Different letters represent significant differences between time points for each treatment (*P* < 0.05).

### Sucrose feeding induces expression of sucrose transporter *SUT2* in the lateral bud

Diverse sucrose transporters are expressed in sink tissues, where they are implicated in a plethora of physiological processes, including seed formation (Weber et al., 1997), tuberization (Kühn et al., 2003) and fruit formation (Davies et al., 1999). We reasoned that a sucrose transporter may be involved in the mobilization of fed sugars from the stem to the bud, and that the expression of that putative transporter might be induced by sugar feeding. To test whether any sucrose transporters are activated during sugar supply, we fed detached etiolated stems with sucrose, hexoses, palatinose or sorbitol and sampled the third lateral bud after 0, 2, 4, 8 and 24 h. The effect of sugar feeding on the gene expression of the three known potato sucrose transporters (SUTs) (Chincinska et al., 2008) was examined.

None of the tested sugars induced a significant change in the expression of *SUT1* or *SUT4* during 24 h of stem feeding (Fig. 3A, C). In contrast, the relative expression of *SUT2* was enhanced 5.8- to 6-fold within 4 h of sucrose feeding, declined after 8 h, and remained at that low level to 24 h. Conversely, *SUT2* transcript level was unaffected by hexoses, palatinose or sorbitol during 24 h of feeding (Fig. 3B). Therefore, the expression of *SUT2* was associated with sucrose translocation into the lateral bud.

**Fig. 3.**
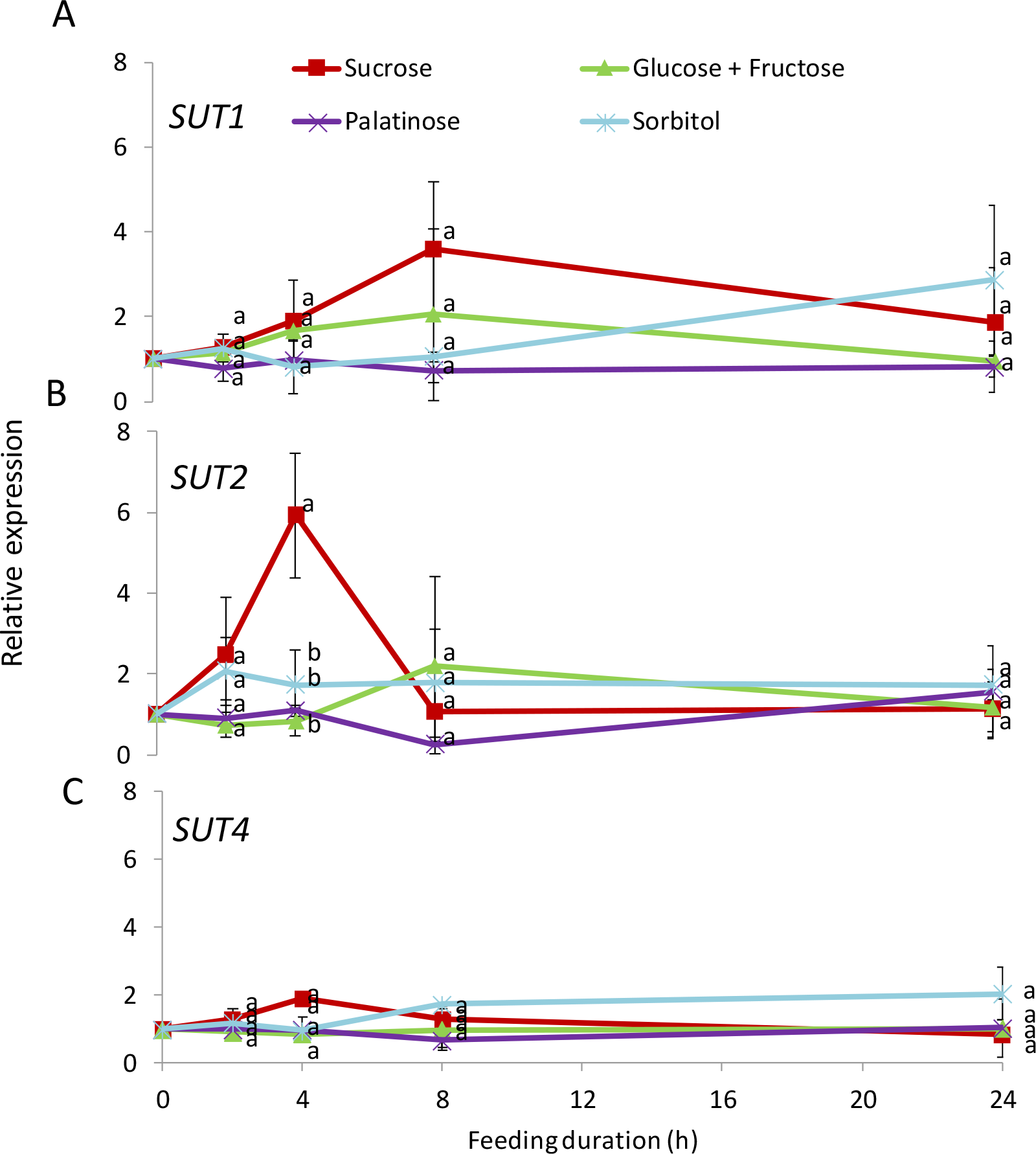
Sucrose feeding of stems induces upregulation of the sucrose transporter *SUT2* in the lateral bud. Detached etiolated stems were fed with 300 mM sucrose, hexoses, palatinose or sorbitol at 14°C, 95% relative humidity, in the dark. The transcript levels of **A**, *SUT1*, **B**, *SUT2* and **C**, *SUT4* were estimated by real-time quantitative PCR. Gene transcript level is expressed relative to controls (0 h) which were set to 1 and normalized to *Elf1* transcript level. Each value is the mean of three independent biological replicates. Error bars represent SE. Different letters represent significant differences between treatments at each time point (*P* < 0.05).

### Sucrose induces the expression and activity of VInv prior to lateral bud elongation

Since hexoses induced stem branching, we hypothesized that bud burst induced by sucrose is mediated by the activity of VInv, a key enzyme involved in sucrose degradation, in the developing bud. To test this hypothesis, etiolated stems were detached and fed with sucrose, hexoses, palatinose or sorbitol for 24 h, and VInv transcript level and activity were determined. After 8 h of sucrose feeding, *VInv* transcript level in the third lateral stem bud was three times higher than its level before feeding, and then decreased back to prefeeding levels after 24 h. VInv activity was enhanced in the lateral bud base (node) of sucrose-fed stems as early as 2 h into feeding, and remained significantly higher until the 24 h measurement. In contrast, VInv transcript level and activity were not induced by other sugars (Fig. 4A, B). These findings provide evidence of a role for VInv in the lateral bud burst induced by sucrose.

**Fig. 4.**
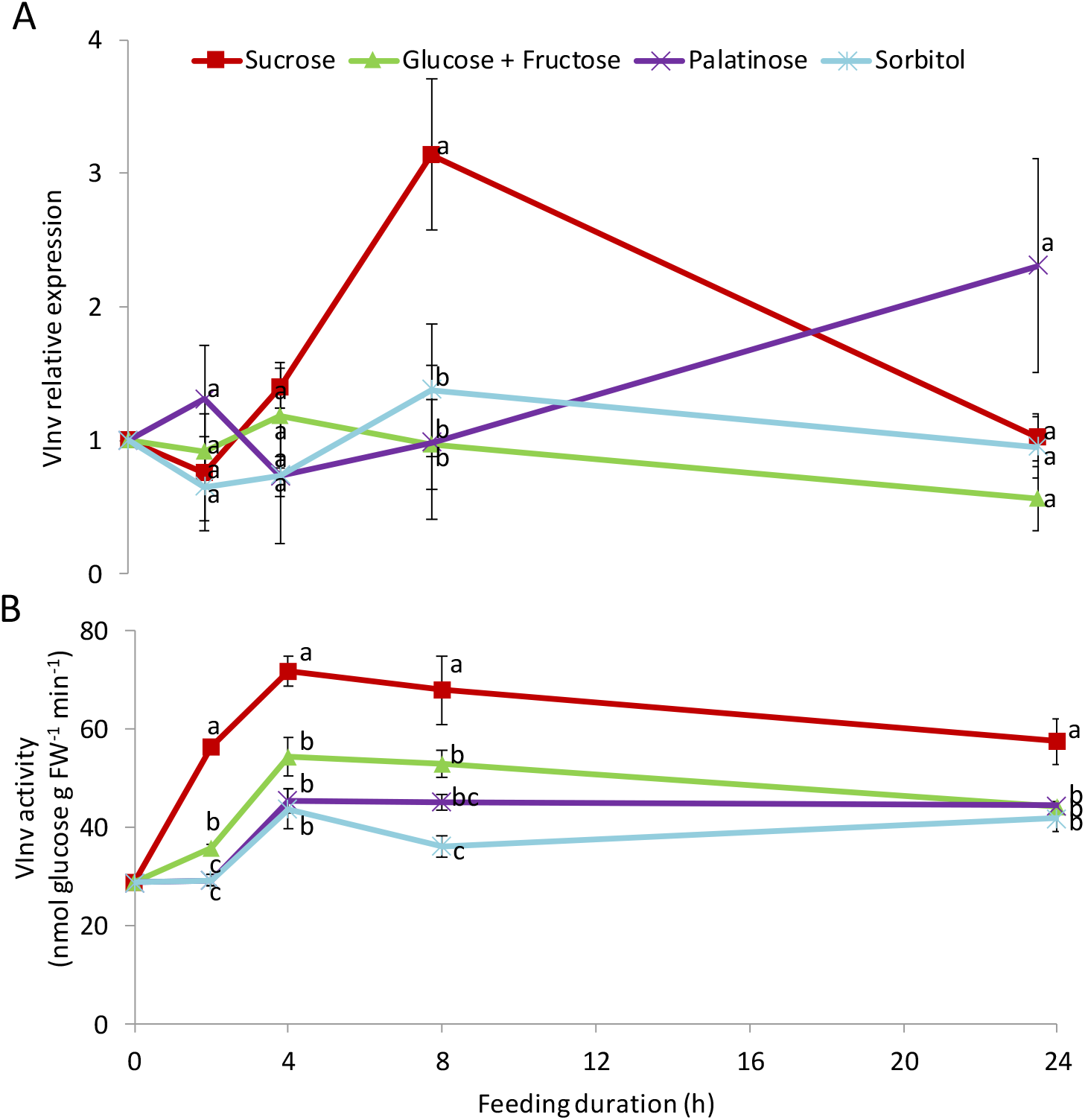
Sucrose feeding of stems induces higher expression and activity of VInv in the lateral bud. Detached etiolated stems were fed with 300 mM sucrose, hexoses, palatinose or sorbitol at 14°C, 95% relative humidity, in the dark. **A**, *VInv* transcript level at the lateral bud was determined by real-time quantitative PCR using gene-specific primers. Gene transcript level is expressed relative to controls (0 h) which were set to 1 and normalized to *Elf1* transcript level. **B**, VInv activity at the stem node. Each value is the mean of three independent biological replicates. Error bars represent SE. Different letters represent significant differences between treatments at each time point (*P* < 0.05).

### VInv is involved in branching

To investigate the involvement of VInv in sucrose-induced stem branching and bud elongation, we compared the effects of sucrose feeding between WT plants and *VInv*-silenced lines. We used four *VInv*-RNA interference (RNAi) lines with a range of *VInv*-silencing levels (Zhu et al., 2014; 2016; Salam et al., 2017). Sucrose-induced branch number was substantially reduced, but not abolished, in *VInv*-RNAi lines compared to the WT, suggesting a role for VInv in the sucrose-induced branching (Fig. 5A). While there was also some effect of the *VInv*-RNAi lines on sucrose-induced lateral bud elongation, it was not statistically significant (Fig. 5B). These results suggest that VInv activity is important for sucrose-induced lateral bud burst but has, at most, a minor role in sugar-induced lateral bud elongation.

**Fig. 5.**
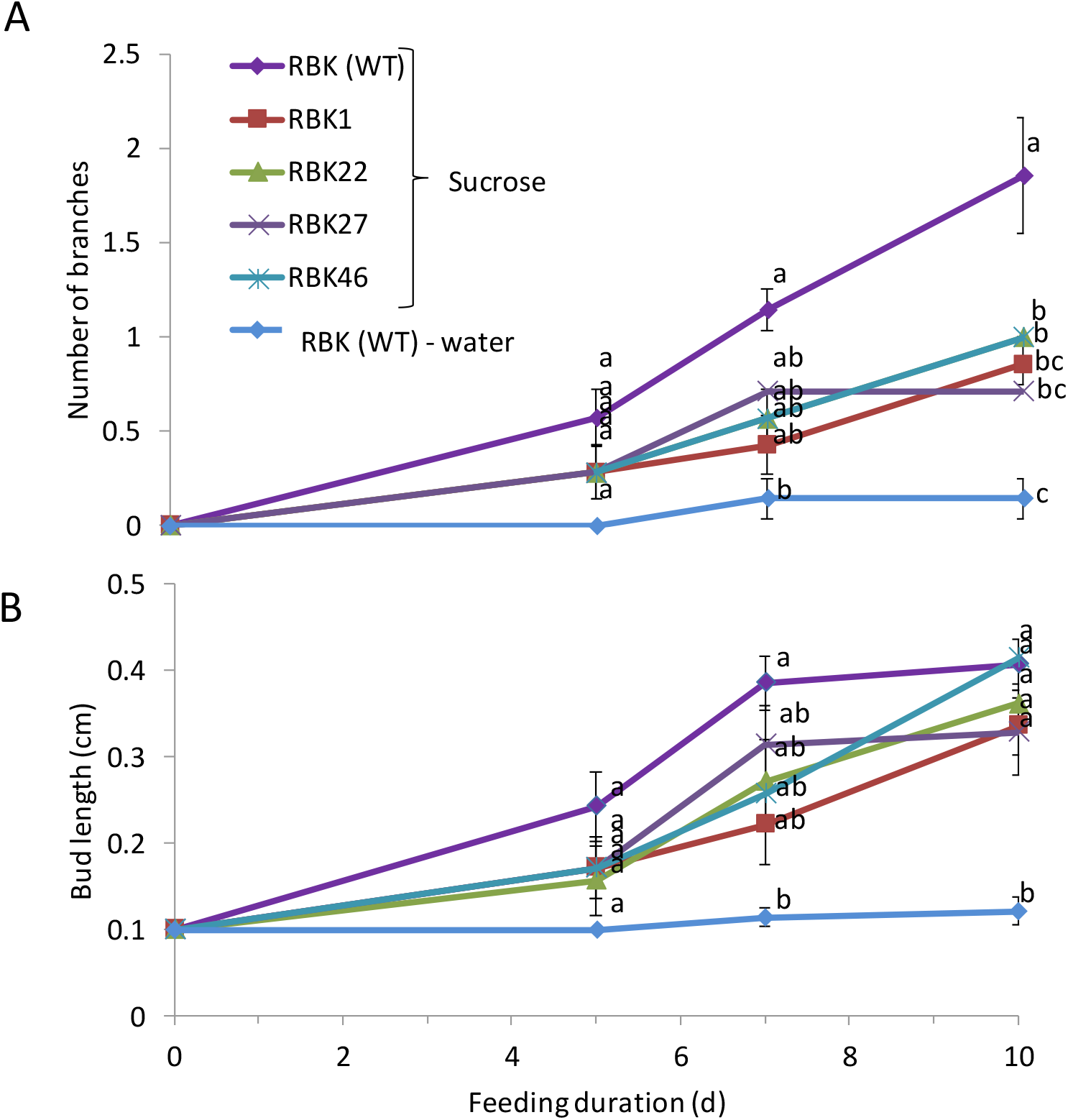
Silencing *VInv* reduces the effect of sucrose on stem branching. Detached etiolated stems of ‘Russet Burbank’ (RBK-WT) were fed with water or 300 mM sucrose, and silenced lines (RBK1, RBK22, RBK27, and RBK46) were fed with 300 mM sucrose for 10 days at 14°C, 95% relative humidity, in the dark. **A**, Number of branches. **B**, Lateral bud length. Data represent averages of two experiments, each performed with seven replicates per treatment. Error bars represent SE. Different letters represent significant differences between treatments at each time point (*P* < 0.05).

### Sucrose triggers accumulation of CK prior to stem-branching initiation

CKs are able to trigger bud outgrowth, and their accumulation often correlates with bud outgrowth in a variety of species, including potato (Bredmose et al., 2005; Shimizu-Sato et al., 2009; Hartmann et al., 2011; Buskila et al., 2016). Moreover, sugars, including sucrose, induce CK synthesis (Barbier et al., 2015b; Kiba et al., 2019). We therefore tested whether CKs are involved in the sucrose-induced bud outgrowth. We quantified their accumulation in the stem node following sugar feeding of etiolated stems. Levels of intermediate (zeatin riboside) and active (zeatin) CK forms increased following feeding with sucrose but not with hexoses or water (Fig. 6), demonstrating that CK accumulation in the lateral bud is induced by sucrose.

**Fig. 6.**
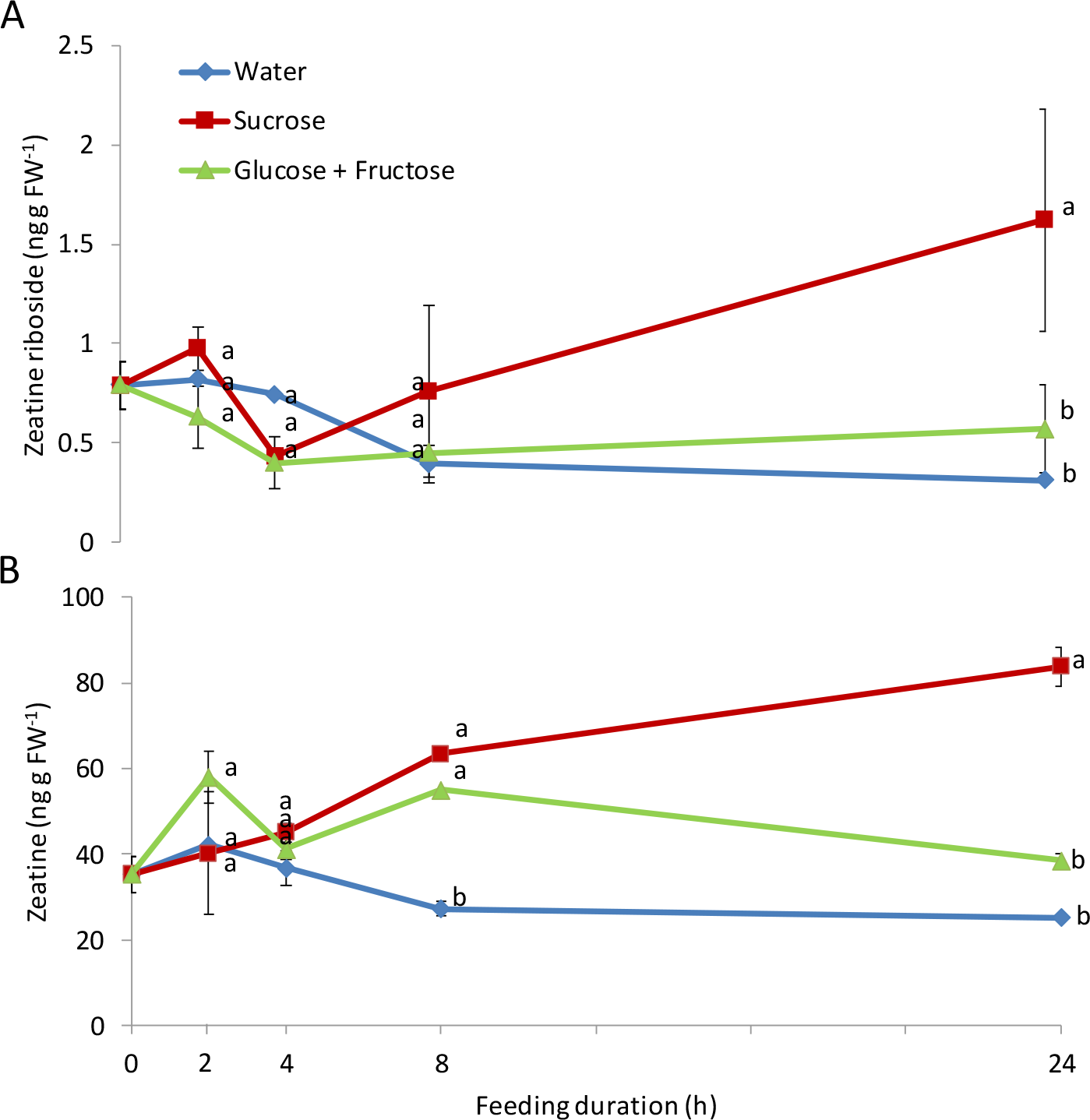
Feeding etiolated stems with sucrose induces higher content of endogenous cytokinin in the node of the lateral bud. Levels of **A**, zeatin riboside and **B**, zeatin in untreated sprouts (0 h) or sprouts supplemented with sugars (sucrose or glucose + fructose at 300 mM) or water, at different time intervals at 14°C, 95% relative humidity, in the dark. Data are means + SE of three measurements. Different letters represent significant differences between treatments at each time point (*P* < 0.05).

Since sucrose caused CK accumulation (Fig. 6), we tested whether exogenous CK application would induce lateral bud outgrowth. Feeding etiolated stems with the synthetic CK, 6-benzylaminopurine (BAP), led to a dose-dependent increase in branching and bud elongation, similar to the effect of sucrose feeding (Fig. 7). Feeding with a mixture of BAP and sucrose significantly increased branching and lateral bud elongation relative to the each treatment alone (Fig. 7).

**Fig. 7.**
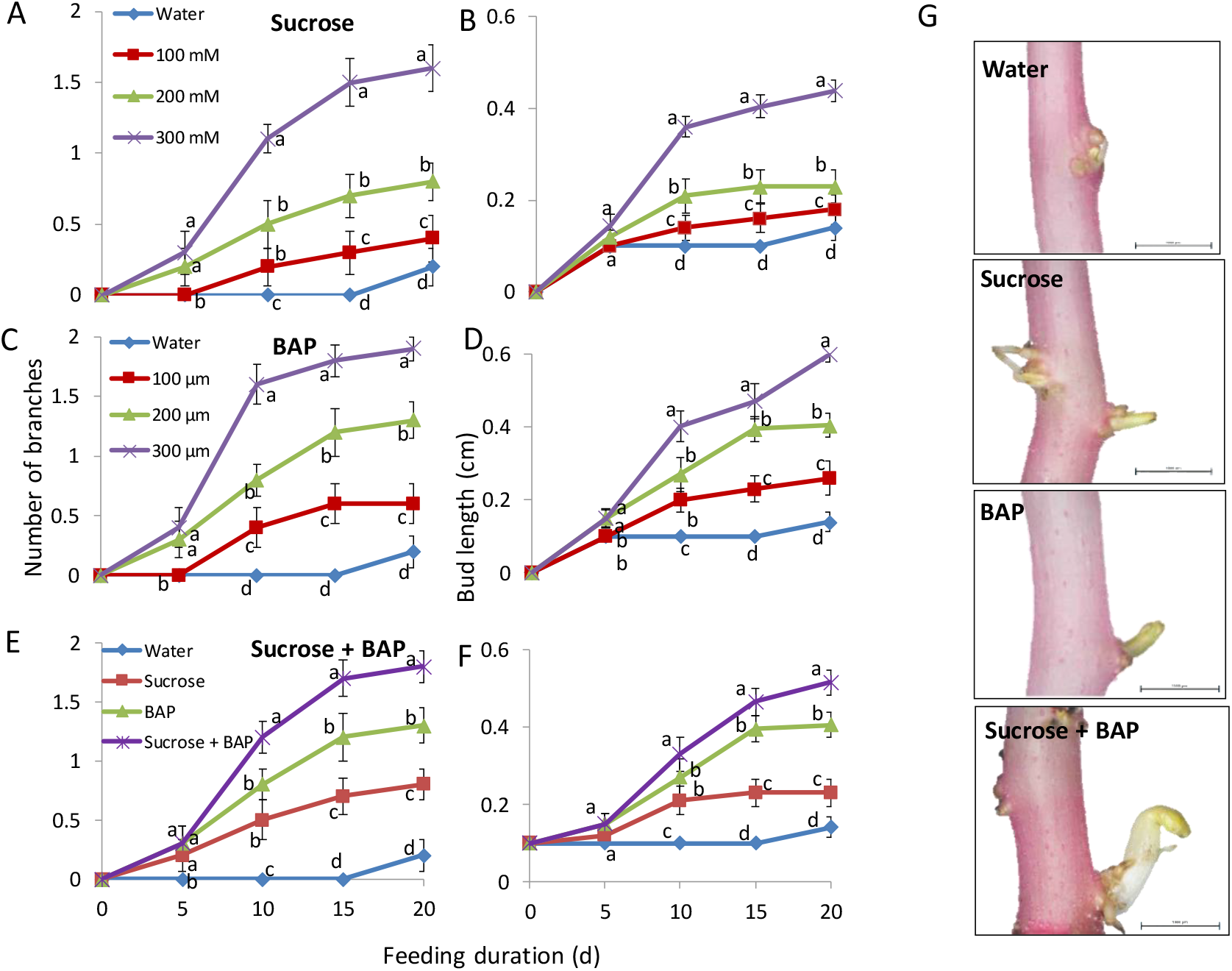
Sucrose and CK have an additive effect on etiolated stem branching and lateral bud elongation. Sprouts were detached from the tubers, incubated at 14°C, 95% relative humidity, in the dark, and fed for 20 days with **A, B**, 0, 100, 200 or 300 mM sucrose; **C, D**, 0, 100, 200 or 300 μm BAP; **E, F**, 200 mM sucrose with or without 200 μm BAP, or water. Number of branches developed and lateral bud length were measured. **G**, Typical lateral bud after 15 days of feeding. Bars = 100 μm. Data represent averages of two experiments, each performed with 10 replicates per treatment. Error bars represent SE. Different letters represent significant differences between treatments at each time point (*P* < 0.05).

To determine whether CK mediates the effect of sucrose on stem branching, we fed inhibitors of CK synthesis (lovastatin) or perception (PI-55, LGR-991) to etiolated stems with sucrose. The effects of sucrose on stem branching and lateral bud elongation were completely suppressed by these inhibitors (Fig. 8). LGR-991 and PI-55 caused repression of bud outgrowth that could not be overcome by BAP application. In contrast, BAP was able to induce bud outgrowth even after feeding with lovastatin (Supplementary Figure S1). This suggests that sucrose requires CK to induce the bud burst and elongation.

**Fig. 8.**
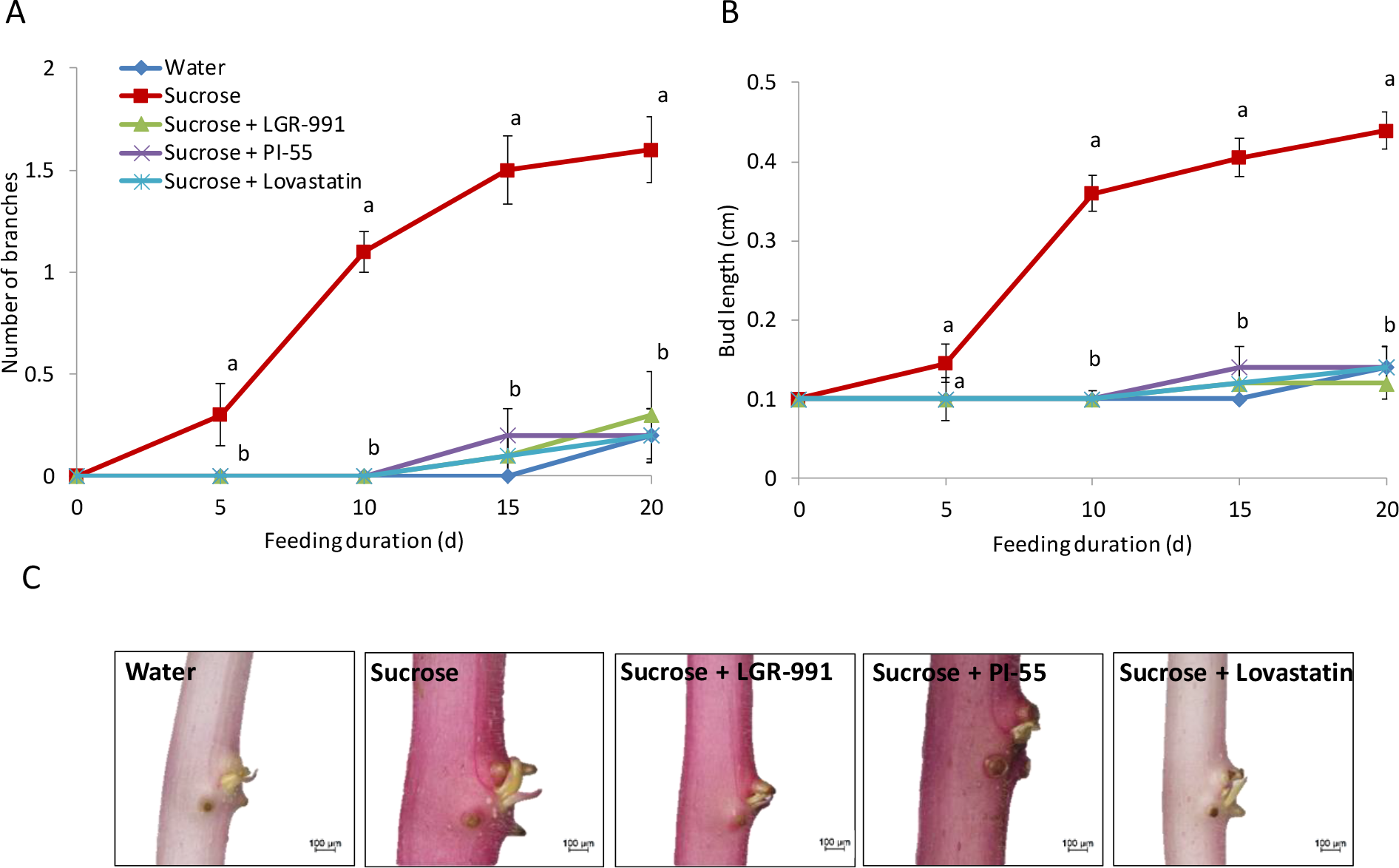
CK inhibitors eliminate branching induction by sucrose. Etiolated stems were detached from the tubers and fed with 300 mM sucrose, 300 mM sucrose with CK-synthesis inhibitor (lovastatin, 200 μm), or with CK-perception inhibitors (LGR-991, Pi-55, 200 μm), or water for 20 days at 14°C, 95% relative humidity, in the dark. **A**, Number of branches. **B**, Lateral bud length. **C**, Typical lateral bud after 15 days. Bars = 100 μm. Data represent averages of 10 replicates per treatment. Error bars represent SE. Different letters represent significant differences between treatments at each time point (*P* < 0.05).

### VInv activity is induced by sucrose and CK

CKs have also been shown to induce VInv expression and nutrient sink strength in bamboo and tobacco (Roitsch and Ehneß, 2000; Werner et al., 2008; Liao et al., 2013). We have shown that glucose and fructose, which are the hydrolytic products of sucrose cleavage by VInv, can induce branching (Salam et al., 2017). However, since a mixture of sucrose and CK inhibitors yielded no branching, we hypothesized that VInv activity is also affected by CK, or that VInv-mediated branching requires CK. To test these hypotheses, we measured the effect of CK inhibitors on sucrose-induced VInv activity. CK inhibitors reduced the effects of both sucrose and BAP on VInv activity (Fig. 9), indicating that sucrose and CK can both trigger VInv activity, and that the impact of sucrose on VInv activity is partially dependent on CK.

**Fig. 9.**
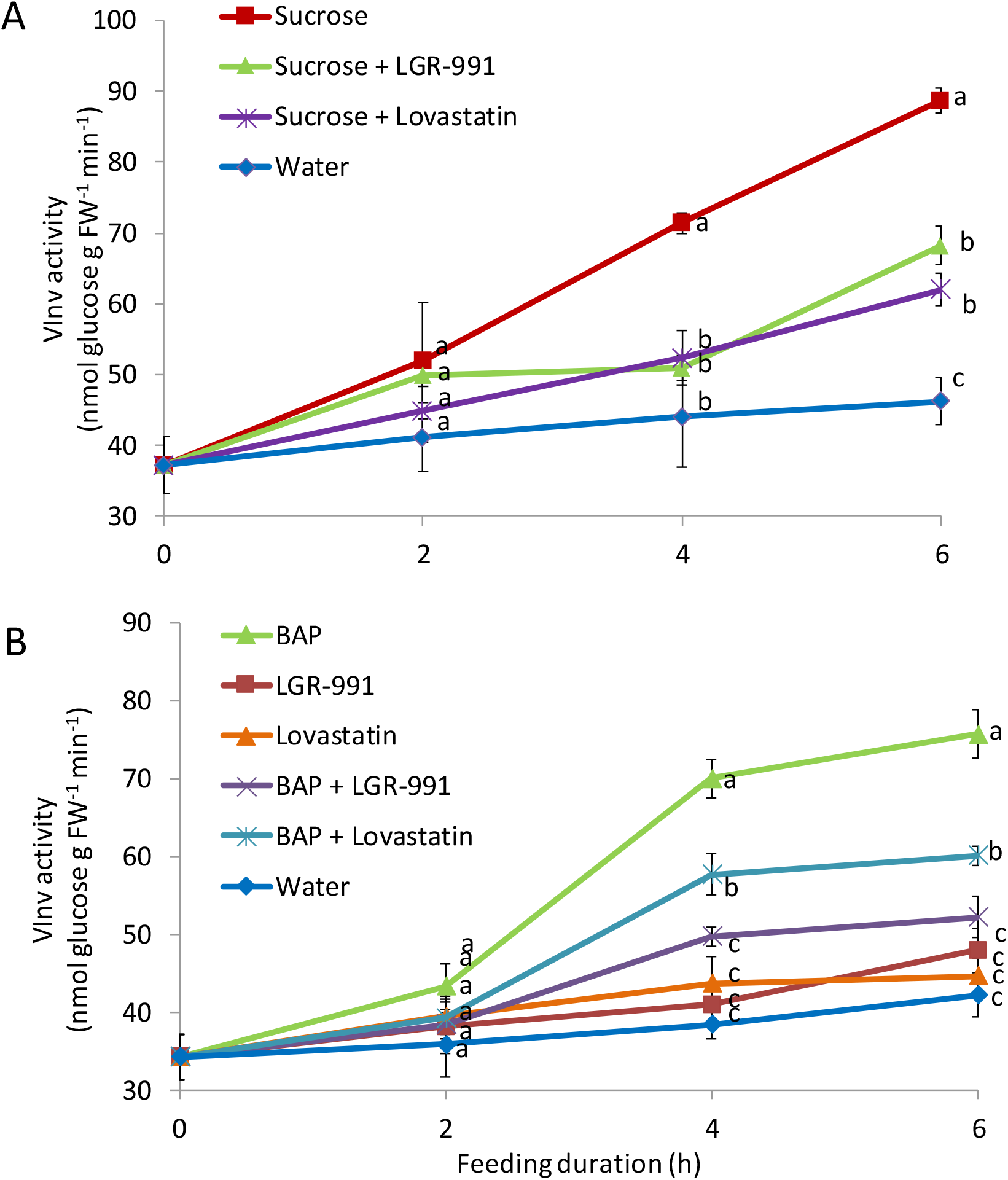
CK inhibitors reduce VInv activity induced by sucrose or BAP. Detached etiolated stems were incubated at 14°C, 95% relative humidity, in the dark and were fed with **A**, 300 mM sucrose, 300 mM sucrose with CK-synthesis inhibitor (200 μm lovastatin) or with CK-perception inhibitor (200 μm LGR-991), or water, **B**, 200 μm BAP, CK-synthesis inhibitor (200 μm lovastatin), CK-perception inhibitor (200 μm LGR-991), BAP with lovastatin or with LGR-991, or water. Data represent averages of five replicates per treatment. Error bars represent SE. Different letters represent significant differences between treatments at each time point (*P* < 0.05).

## DISCUSSION

### Sucrose moves into the lateral bud to induce its burst and elongation

Sugars play a major role in plant growth and development; they provide energy and a source of carbon for protein and cell-wall synthesis (Patrick et al., 2013). Independent of their nutritional role, sugars also play a signaling role and can therefore interact with other regulatory networks to control plant development (Lastdrager et al., 2014; Yadav et al., 2014; Li and Sheen, 2016; Sakr et al., 2018). Recent studies suggested that sugars are important signaling regulators of bud outgrowth. Using an elegant set of experiments, Mason et al. (2014) demonstrated the systemic movement of sucrose as a branching signal from the leaf to the lateral bud after decapitation. Barbier et al. (2015a) demonstrated that sugar availability, together with auxin (Bertheloot et al., 2019), control the entrance of buds into sustained growth. Recently, we showed that sucrose and its hydrolytic products can induce stem branching in a dose-responsive manner under etiolated conditions (Salam et al., 2017). Taken together, these results suggest that bud branching and elongation are closely linked to the mobilization of sugars toward the buds. Here, feeding sprouts exogenously showed a differential effect of sucrose and hexoses on the number of branches and lateral bud elongation (Fig. 1). Treatment with water, a slowly metabolizable analog of sucrose (palatinose), or the osmotic control sorbitol did not induce significant VInv activation or lateral bud growth (Figs 1 and 4). Palatinose is an isomer of sucrose that differs in its glyosidic linkage between glucose and fructose. Palatinose was neither cleaved nor taken up by tomato cells in a suspension culture (Sinha et al., 2002). In sugarcane cells, palatinose is not actively transported in the cell and can only be partially cleaved (10% compared to sucrose) by vacuolar invertases and sucrose synthase when the cells are damaged (Wu and Birch, 2011). In addition, our data support previous findings of lack of recognition or transportation of turanose and palatinose by sucrose transporters (Z.-S. Li et al., 1994; M’Batchi & Delrot, 1988). The lack of cellular transport and VInv cleavage may explain why palatinose did not initiate branching and elongation in our system. Taken together, our results are consistent with previous reports of sucrose’s central role in bud release (reviewed in Barbier et al., 2019). The lower effect of sucrose-degradation products suggests that sucrose not only acts as an energy source, but also through other pathways, to initiate branching.

In etiolated stems in the dark, sugars are expected to move through the stem mainly to the strongest sink—the apical bud (Buskila et al., 2016). In our system, feeding the etiolated stem with exogenous sucrose activated the expression of *SUT2,* a specific sucrose transporter, inside the lateral bud tissue. Previous studies have reported and emphasized the importance of sucrose transporters in various sink tissues, and their intricate roles in diverse physiological processes, such as flowering (Chincinska et al., 2008), latex synthesis (Dusotoit-Coucaud et al., 2009), pollen development (Lemoine et al., 1999; Takeda et al., 2001), and tuberization (Chincinska et al., 2008). Similar to our findings, Henry et al. (2011) demonstrated in *Rosa* that three of the four putative sucrose transporters (*RhSUC2*, *RhSUC3* and *RhSUC4*) are expressed in the bud. They concluded that only *RhSUC2* expression is correlated with bud break in decapitated plants, and that it plays a central role in sucrose influx into outgrowing buds. Chincinska et al. (2008) reported the prominence of *SUT2* expression in sink tissues of potato tubers. Decourteix et al. (2008) demonstrated that in walnut stems, expression of *JrSUT1* is correlated to increasing bud sink strength during outgrowth. Furthermore, using radiolabeled sugars, we demonstrated that the sugar is imported into the lateral bud during bud outgrowth (Fig. 2). These modifications are in accordance with an increase in plasmalemma ATPase activity (Aue et al., 1999; Alves et al., 2001; Alves et al., 2007) and active sugar absorption (Marquat et al., 1999; Maurel et al., 2004b; Lecourieux et., 2010) in the bud or neighboring stem region. Our data are in line with these previous reports and clearly show that the lateral buds become stronger sink organs upon sugar feeding.

### Sucrose cleavage to hexoses is required for stem branching

Sucrose feeding of etiolated stems induces a sequential transcript accumulation of *SUT2,* peaking at 4 h (Fig. 3); this is followed by increased expression of *VInv* at 6–8 h of sucrose feeding (Fig. 4). VInv has been shown to be a major component of organ sink strength (Nägele et al., 2010; Albacete et al., 2015) and cell elongation (Morris and Arthur, 1984; Morey et al., 2018). The imported sucrose can contribute to cellular growth processes by contributing to the carbon skeleton and energy, and by providing osmotically active molecules for cell expansion. Similarly, the sucrose imported into the bud, or its cleavage products (hexoses) derived from the action of *VInv*, may serve as signal molecules to regulate genes involved in development (Li and Sheen, 2016; Wang et al., 2018; Gibson, 2005; Barbier et al., 2019).

Silencing of *VInv* results in inhibition of sucrose cleavage (Bhaskar et al., 2010; Zhu et al., 2016). Salam et al. (2017) showed that a higher sucrose level was correlated with higher branching of transgenic potato tubers. Here, feeding sucrose to stems of RNAi lines with different levels of *VInv* silencing revealed significantly lower numbers of branches but no significant effect on bud length (Fig. 5). Additionally, palatinose, which is only poorly cleaved into hexoses in this system, could not trigger bud outgrowth. These results support the role of hexoses, produced by VInv in the lateral bud, in stem-branching induction. Heyer et al. (2004) overexpressed cell wall or cytosolic invertase in *Arabidopsis*, and this led to changes in the shoot-branching pattern in a composite manner, differentially affecting the formation of axillary inflorescences, branching of the main inflorescence, and branching of side inflorescences. The essential role of acid invertases in regulating sink strength was analyzed in transgenic carrot plants by antisense suppression of *VInv* under control of the 35S-CaMV promoter that is predominantly active in carrot tap roots. The resulting lowered carbohydrate content in the roots and severe impairment of both growth and development demonstrated the important function of VInv in sucrose partitioning (Tang et al., 1999). In addition to modulating sink strength, antisense suppression of *VInv* in tomato led to differential growth, and alterations in fruit size (Klann et al., 1996). Goetz et al. (2001) reported that antisense repression of the cell wall invertase *Nin88* results in assimilate blockage and developmental arrest during early stages of pollen development, leading to a distorted and invaginated morphology. The transgenic lines revealed a correlation between reduced enzymatic activity and decreased germination efficiency. Exogenous supply of glucose or sucrose partly rescued developmental arrest (Mārc Goetz & Roitsch, 2006), suggesting that the function of invertase is not only to provide carbohydrates to sustain growth, but also to create a delicate, fine-tuned balance between the sucrose and hexose sugars required as metabolic signals to regulate growth and development. It may also suggest a role for VInv as a sugar sensor. Indeed, enzymes catalyzing sugars, such as Hexokinase1 or Fructose-1,6-bisphosphatase, have been shown to play a sensor role in sugar signaling (Cho & Yoo, 2011; Moore et al., 2003). It would be interesting to test whether VInv is a sensor for the sucrose pathway during bud outgrowth.

### Sucrose promotes CK accumulation in the lateral bud

CKs are known to promote bud release from dormancy in intact plants (Sachs and Thimann, 1964; 1967; Dun et al.ArromArroma, 2012; Kalousek et al., 2014). However, how they are induced to accumulate during bud outgrowth remains unclear. Our results demonstrate that sucrose upregulates CK accumulation in stem nodes (Fig. 6). Compared to hexoses and water, sucrose induced the accumulation of intermediate and active forms of CKs in the bud node prior to lateral bud burst. Feeding with a mixture of BAP and sucrose increased the effect over that of each component alone with respect to both branching level and lateral bud elongation (Fig. 7). The effect of sugars on CK production has been reported for Lily flowers (Arrom & Munné-Bosch, 2012) and *Arabidopsis* seedlings (Kushwah and Laxmi, 2014; Kiba et al., 2019). Sucrose has been reported to strongly induce CK synthesis in *in vitro*-grown single nodes of rose in absence of auxin, suggesting that CKs might mediate the effect of sucrose, although the authors concluded that CKs alone are not sufficient to stimulate bud outgrowth in *Rosa* single nodes (F. Barbier et al., 2015). In presence of auxin in the growth medium, sucrose could not promote CK accumulation, and the CK content did not correlate with the onset of bud outgrowth (Bertheloot et al., 2019). Here, we report here that CKs play an important role in mediating sucrose-promoted bud outgrowth in etiolated potato stem. Indeed, sucrose-induced bud outgrowth was significantly suppressed by inhibitors of CK synthesis or perception (Fig. 8).

CKs have been reported to enhance sugar sink strength in the tissues in which they accumulate, notably through upregulation of invertases (Fig. 9) and cell-cycle promotion (Roitsch and Ehneß, 2000; Peleg et al., 2011 Wang et al., 2016). Similar to our findings, Wang et al. (2016) reported that when *tasg1* mutants were treated with the CK inhibitor lovastatin, the activity of invertase was inhibited, and this was associated with a premature senescence phenotype. However, the activity of invertase was partially recovered in *tasg1* treated with BAP, suggesting that CKs might regulate the invertase activity involved in sucrose remobilization. Our findings are consistent with previous studies in which CKs were shown to adjust the sugar partitioning and sink strength of some organs through the regulation of sugar transporters and invertases ((Thomas, 1986; Roitsch and Ehneß, 2000; Guivarc’h et al., 2002; Werner et al., 2008; Proels and Roitsch, 2009; Liao et al., 2013). In addition, our results are in agreement with recent results obtained by Roman et al. (2016) in rose buds showing that exogenous feeding of CK induces SUC2 and VInv, although an environmental cue— light—was integral to their system. Taken together, our data strongly suggest a crucial role for CKs in the sucrose-induced axillary bud outgrowth in etiolated stem through increased VInv activity in the bud, leading to a possible increase in sink strength.

In summary, our study demonstrates that sucrose induces bud growth and elongation better than its moieties or poorly cleavable analog palatinose. Sucrose activates CK accumulation, whereas hexoses do not affect CK levels. Sucrose and CK induce higher VInv activity, which contributes to increase the nutrient sink strength required to promote bud outgrowth. The induced activity of VInv in the lateral bud leads to sucrose degradation to hexoses, providing energy and a distinct profile of sugar signals that can have profound developmental effects on lateral bud growth (Fig. 10).

**Fig. 10.**
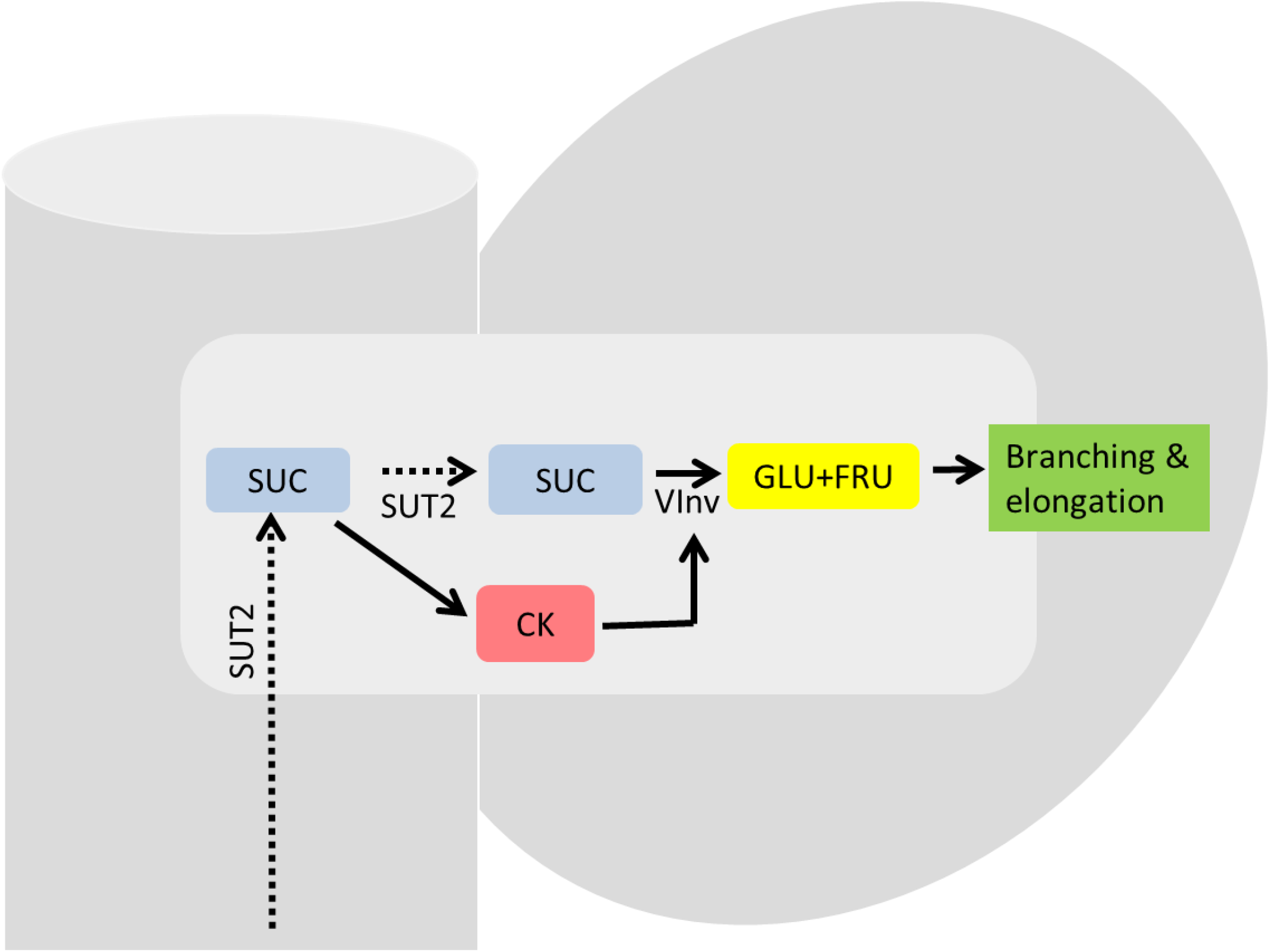
Schematic model for the impact of sucrose on stem branching and lateral bud elongation. Sucrose (SUC) is transported by SUT2 into the lateral bud. Sucrose availability in the lateral bud triggers the synthesis of CK. CK induces VInv activity. VInv degrades sucrose to its hydrolytic products (GLU+FRU). Hexoses in the lateral bud support branching and elongation. Dotted arrows represent hypothetical interactions.

## ACKNOLEGMENTS

This research was supported by BARD (U.S.–Israel Binational Agricultural Research and Development fund) project IS-5038-17C.

---

**Supplementary Table S1.**
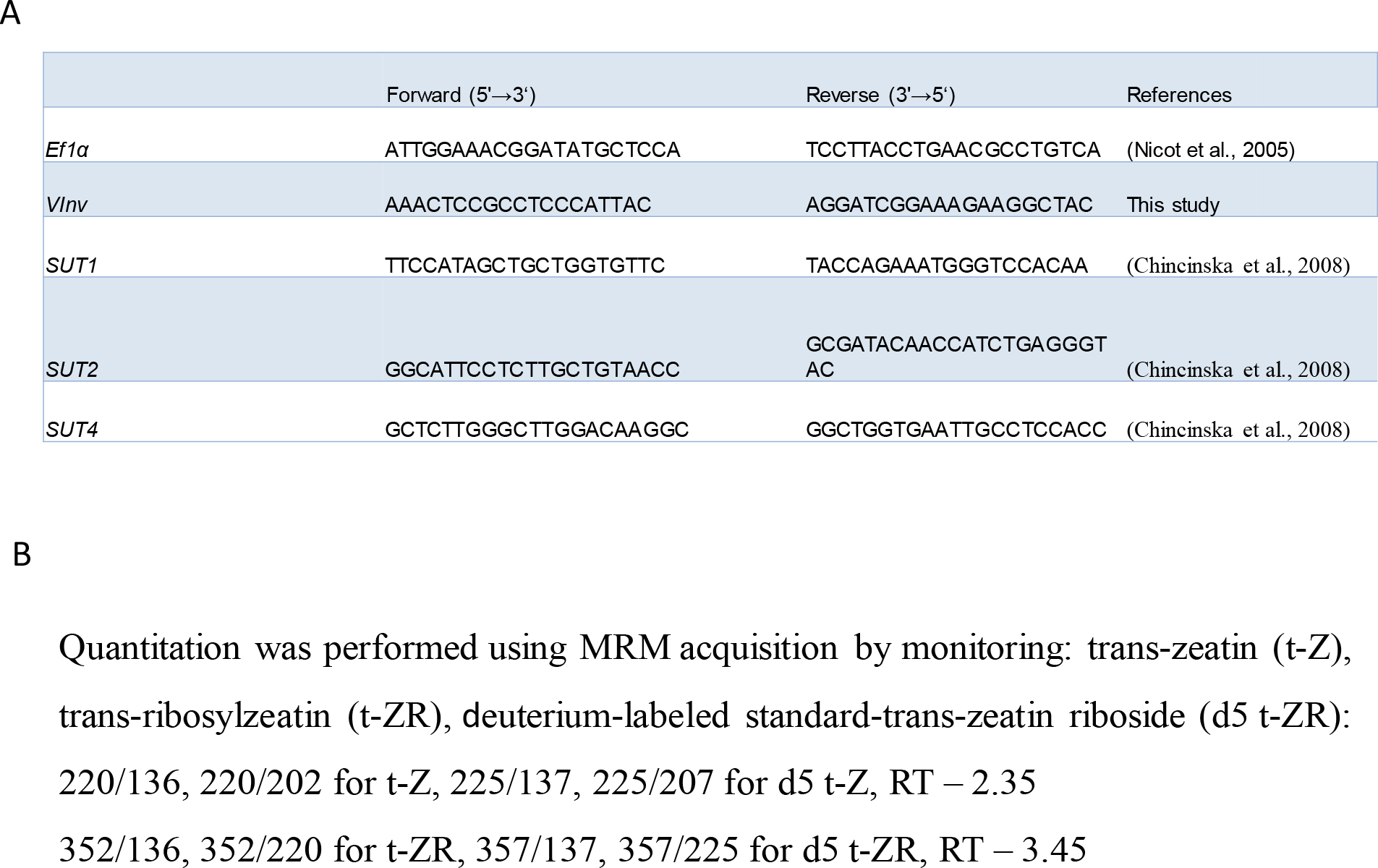
**A**, Primers used for qRT-PCR in this study. **B**, MRM data-acquisition parameters for hormones.

**Supplementary Fig. S1.**
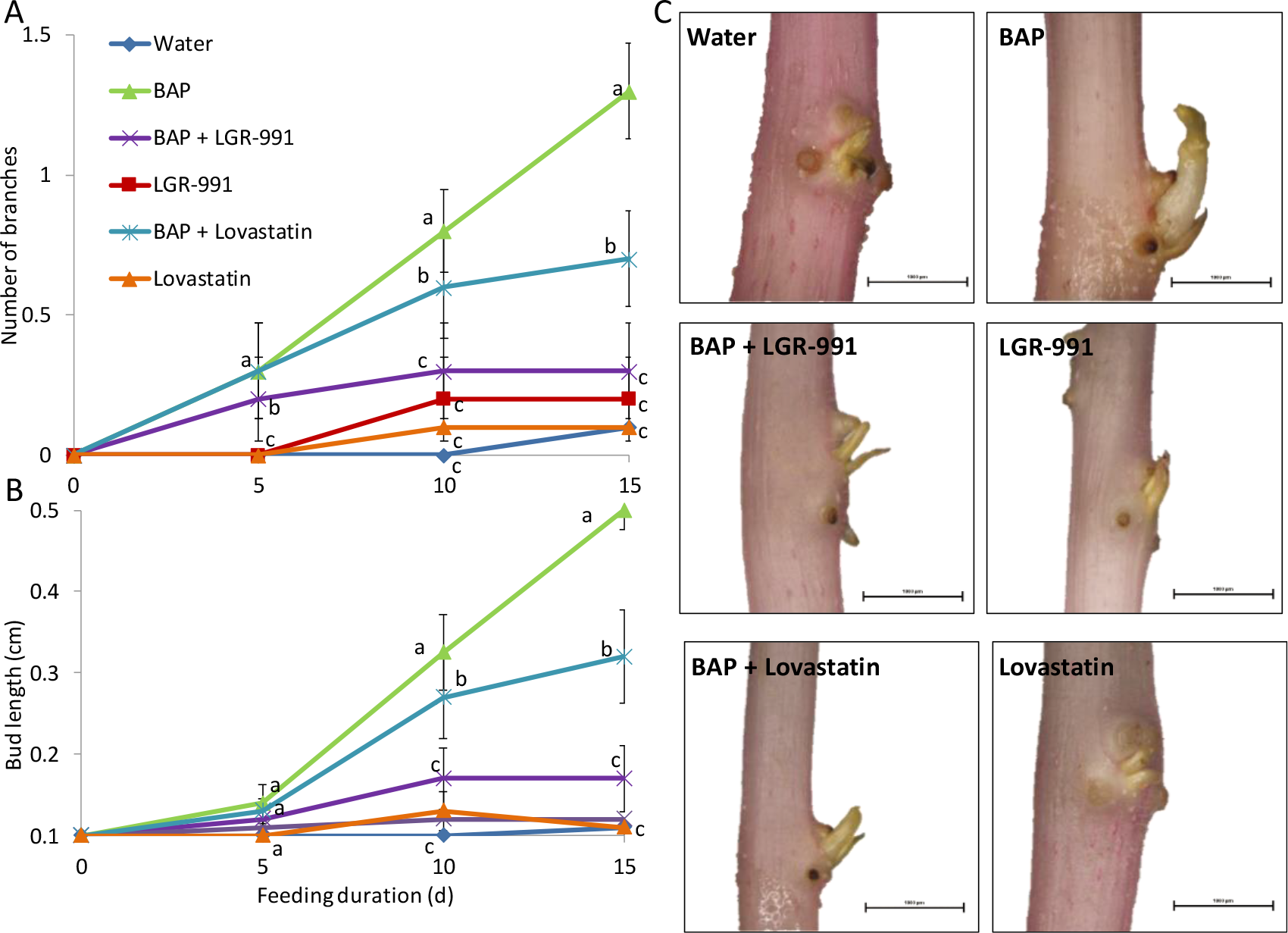
Effects of cytokinin (CK), CK inhibitors, or a mixture of CK and its inhibitors on bud outgrowth and elongation. Sprouts were detached from the tubers and supplemented with a synthetic form of CK (6-benzylaminopurine, BAP, 200 μm), CK-synthesis inhibitor (lovastatin, 200 μm), CK-perception inhibitor (LGR-991, 200 μm), BAP with lovastatin or with LGR-991, or water for 10 days at 14°C, 95% relative humidity, in the dark. **A**, Number of branches and **B**, bud lengthwere measured for 15 days of treatment. **C**, Images showing sprouts with or without branches after 10 days of treatment. Bars = 100 μm. Data represent averages of 10 replicates per treatment. Error bars represent SE. Different letters represent significant differences between treatments at each time point (*P* < 0.05).

